# Repeated Inertial Stimulation Reveals Divergent Plasticity in Peripheral and Central Vestibular Pathways

**DOI:** 10.64898/2026.09.22.752866

**Authors:** Syed Danial Naqvi, Mamiko Niwa, Rod D. Braun, Mirabela Hali, Aaron K. Apawu, Avril Genene Holt

## Abstract

The vestibular system consists of peripheral end organs that provide the brain with temporally precise information about head position in space required for motor coordination and postural stability. Irregular vestibular afferents of the inner ear are particularly sensitive to rapid changes in acceleration, or jerk, which is necessary for making quick, corrective movements and postural adjustments. Vestibular short-latency evoked potentials (VsEPs) are waveforms that provide a measure of synchronous irregular afferent activity originating from peripheral (early peaks) and central (later peaks) vestibular pathways. These VsEPs are elicited by head jerk stimuli, commonly used preclinically. Previous studies have shown that early VsEP waves are dependent upon jerks of an adequate intensity, number, and duration. However, less is known about jerk characteristics necessary for generation of reliable VsEP signals centrally (later peaks). The present study examines the precise temporal dynamics of peripheral and central vestibular centers across a broad range of mechanical vestibular load in rats. We found differential changes in synchrony and timing driven by jerk repetition, intensity, and duration between early and late VsEP waves representing distinct anatomical components of the vestibular circuit. Our results suggest that central components are uniquely sensitive to jerk duration, intensity, and repetition, indicating a disconnect between peripheral and central neuronal recruitment, adaptive plasticity, and time-frequency dynamics. This work provides a foundation for using VsEP to assess jerk-induced physiological vestibular dysfunction and to understand how central relays differentially adapt to variations of mechanical vestibular stress compared with the precise, high-fidelity transmission observed in the periphery.

## 1. INTRODUCTION

The mammalian vestibular system functions as a sophisticated inertial guidance apparatus, utilizing a hierarchy of peripheral sensors and central processing units to maintain postural equilibrium and gaze stability during rapid head movements. Within the otolith organs, the utricle and saccule, this sensory task is divided between distinct afferent populations. While regular afferents are optimized for encoding sustained head orientation, the irregular afferents of the striolar region exhibit a specialized sensitivity to rapid changes in acceleration, or kinematic jerk 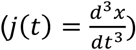 [1-3]. This precise sensitivity is fundamentally rooted in the mechanical properties of the otoconial sensory apparatus, enabling the sub-millisecond temporal precision required to drive the fastest reflex pathways in the body [4].

The vestibular short-latency evoked potential (VsEP) is widely used in preclinical settings as a direct temporal measure of this synchronous neuronal activity [5-7]. Foundational work has firmly established that the initial waves of the VsEP depend strictly on jerk levels rather than peak acceleration [5, 8]. However, previous physiological investigations have primarily operated within a narrow, moderate range of stimulus intensities (<5.5 g/ms) and jerk durations [9]. Furthermore, standard methodologies inherently rely on averaging multiple, repetitive jerk presentations to extract a viable signal. As a result, the physiological limits of this high-fidelity signaling system, specifically the temporal dynamics and threshold behaviors under extreme or highly repetitive inertial loads, remain almost entirely unexplored.

Current electrophysiological models indicate that the initial wave of the VsEP waveform (P1) represents the compound action potential of peripheral irregular afferents [10], whereas subsequent waves (P2 and P3) represent central vestibular responses within the vestibular nuclear complex (VNC) and subsequent central relays [11]. We hypothesized that while the vestibular periphery acts as a robust, high-fidelity transient detector limited by finite structural recruitment, central vestibular pathways process extreme inertial stimuli in a highly state-dependent manner.

To test the functional limits of central vestibular plasticity, the present study systematically examines the effects of varying linear jerk stimulus intensity, duration, and massive repetition on the distinct components of the VsEP. By expanding the stimulus range to extreme intensities (up to 8.8 g/ms) and applying trial-by-trial continuous wavelet transform (CWT) analyses, we reveal a profound dissociation in dynamic range between the stability of the vestibular periphery and the adaptability of central networks

## 2. METHODS

### 2.1. Subjects and design

All procedures were approved by the Institutional Animal Care and Use Committee (IACUC) at Wayne State University. Adult male Sprague-Dawley rats (age 8 – 10 weeks) were obtained from Charles River Laboratory. All rats were individually housed and maintained at 72°F with a 12–hour light/dark cycle. Standard housing conditions with free access to normal rat chow and tap water were provided. Wayne State University (WSU) maintains AAALAC accredited animal facilities under the Division of Laboratory Animal Resources (DLAR). After arrival at the WSU DLAR facility, animals were allowed to acclimate for at least 48 hours before undergoing any procedures.

Rats used to assess the effects of jerk intensity and repetition were randomly assigned to one of five stimulation groups: sham (non-stimulated), 0.64, 3.3, 5.2, or 8.8 g/ms. In contrast, VsEP responses across jerk durations were compared within the same animal using a separate cohort of rats. All rats underwent surgery, as described below, to secure a nut to the skull for attachment to the mechanical shaker.

### 2.2. Surgical procedures

All surgical procedures were performed under aseptic conditions. Animals were anesthetized with ketamine (75 mg/kg) and xylazine (8 mg/kg) administered IP. After stabilization and leveling of the head using a stereotaxic frame, bregma and lambda were exposed, and two anchor screws were placed into the skull. A 6-32 HEX M/S ceramic nut (Small Parts Inc., model #B000FN0BZG, UNPSC code 31161700) or a custom-made titanium head bolt was centered on bregma and affixed to the skull using C&B Metabond cement (Parkell, Edgewood, NY). The ceramic nut or head bolt was further secured to the skull using dental acrylic (Lang Dental Manufacturing Co., Inc. Wheeling, IL). The animals were given 7-10 days to recover prior to VsEP experiments.

### 2.3. Jerk stimulation and vestibular short-latency evoked potential (VsEP)

For jerk stimulation/VsEP testing, animals were anesthetized with an intramuscular injection of ketamine (75 mg/kg) and xylazine (8 mg/kg). Needle electrodes (23 gauge, BD (Becton, Dickinson and Company) were positioned on the vertex of the skull (active), right ear (reference), and hip (ground). The electrode signals were filtered, digitized, and amplified using Spike2 software and CED Power 1401 data acquisition system. Custom Spike2 scripts were used to drive the stimulus and collect VsEP and jerk information. Signals were analyzed using custom MATLAB (MathWorks, Natick, R2019a). VsEP amplitudes were calculated as the difference between the peak and trough voltages (e.g., P1 – N1, P2 – N2). Latencies were calculated as the time of each peak occurred relative to the peak of the jerk stimulus.

Animals were placed in a supine position and the head was attached to a mechanical shaker arm (via ceramic nut) fitted with an accelerometer (Fig. 1A). Rats were exposed to a total of 15 trials at one jerk intensity (0.64, 3.3, 5.2, 8.8 g/s) split across three blocks comprising five trials each (Fig. 1B). A series of 200 jerks (100 up and 100 down) was designated as a trial. A 10-minute rest period was introduced after each block. Animals used for assessing the impact of jerk duration were similarly exposed to a total of 15 trials at one intensity (3.1 g/ms), divided into three blocks corresponding to different durations (0.95, 1.20, or 1.45 ms) presented in a randomized order.

**Figure 1.**
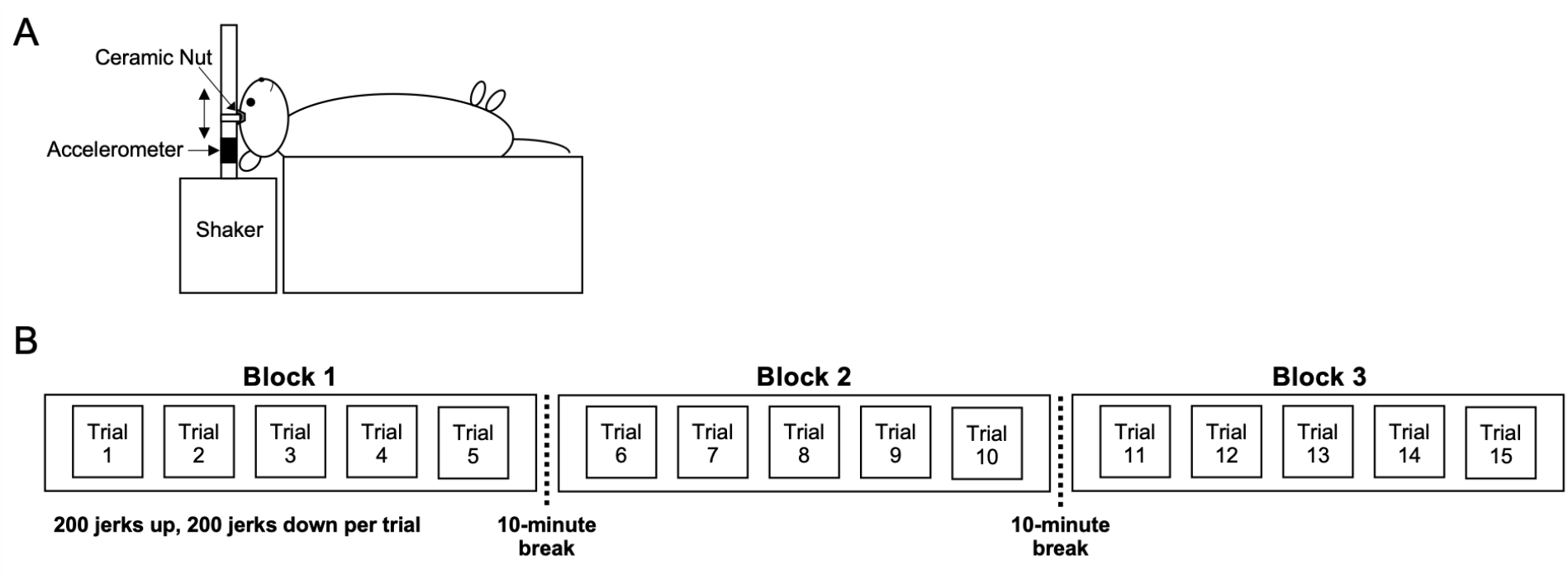
Experimental timeline and jerk stimulation paradigm. A ceramic nut affixed to the skull was attached to the shaker arm in the orientation as shown. Jerk stimuli were applied in the naso-occipital plane (double-headed arrow). All rats were placed supine on the shaker platform **(A)**. The jerk stimulation paradigm included a 10-min quiescent interval between each block. Each block consisted of five trials with 400 bidirectional jerks delivered in each trial, 2,000 jerks per block, for a total of 6,000 jerks. All jerk stimulated rats were exposed to a total of 15 trials, split into three blocks of five trials each **(B)**. Jerk intensity for intensity/repetition experiments was 0.64, 3.3, 5.2, or 8.8 g/s. Jerk duration experiments used a constant intensity of 3.1 g/ms with each block representing a jerk duration of either 0.95, 1.20, or 1.45 ms (randomized order across animals).

### 2.4. Wavelet transform

A continuous wavelet transform with Generalized Morse wavelets was applied to single-trial VsEP traces [12]. Here, a pair of positive- and negative-jerk responses were averaged, and the average response was considered as a single-trial response. Averaging positive- and negative-jerk responses cancel large, stimulus-locked waves in opposite directions, which are thought as an artifact generated by the electro-magnetic shaker that produces the jerk stimuli. Each single-trial VsEP trace encompassed 6 ms before and after the onset of a jerk, having a total of 12 ms length.

The Wavelet Toolbox from MATLAB R2022b (Mathworks, MA) was used to produce the continuous wavelet transform of single-trial VsEP traces. We used a continuous Morse wavelet transform (MATLAB function ‘cwt’) with a symmetry parameter, gamma (γ), at 3 (symmetrical in time domain), a time-bandwidth product, P, at 10 (the square root of P is proportional to the wavelet duration in time), and a redundancy of 10 voices per octave (frequency points per octave in wavelet transform), applied on single-trial VsEP traces. The time-bandwidth product was selected at 10 to shorten the temporal duration of the wavelets (to have better localization in time) as much as possible while minimizing loss of frequency selectivity.

Inter-trial-phase-coherence (ITPC): The phase of the WT coefficient was first determined for every single-trial trace of the VsEP using the MATLAB function ‘angle’. Then, ITPC was determined as: The phase of the WT coefficient was first determined for every single-trial trace of the VsEP using the MATLAB function ‘angle’. Then, ITPC was determined as:

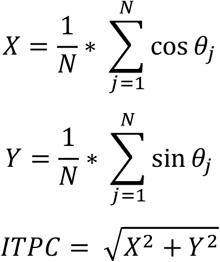

θ_j_ is the phase of the WT coefficient for the j-th single-trial trace of the VsEP and N is the number of trials.

### 2.5. Statistical analysis

All data analysis was performed using Prism 10 software (GraphPad) or MATLAB R2022b. Statistical tests used for each analysis are described in the results section. Statistical significance was defined as p < 0.05.

## 3. RESULTS

### 3.1. Late-stage central vestibular components are uniquely sensitive to jerk stimulus duration

To isolate the temporal processing capabilities of the vestibular pathway, we first investigated te impact of varying jerk stimulus duration (0.95, 1.20, and 1.45 ms) while maintaining a constant intensity of 3.1 g/ms. Evaluation of the resulting VsEP waveforms revealed that the duration of the kinematic stimulus selectively impacts late-stage central integrations, leaving peripheral and early central responses invariant. Specifically, the amplitudes and latencies of the P1 and P2 components, reflecting the compound action potentials of the irregular afferents and early vestibular nuclear complex (VNC) activity, respectively, were unaffected by changes in stimulus duration (Fig. 2A-B, D-E).

**Figure 2.**
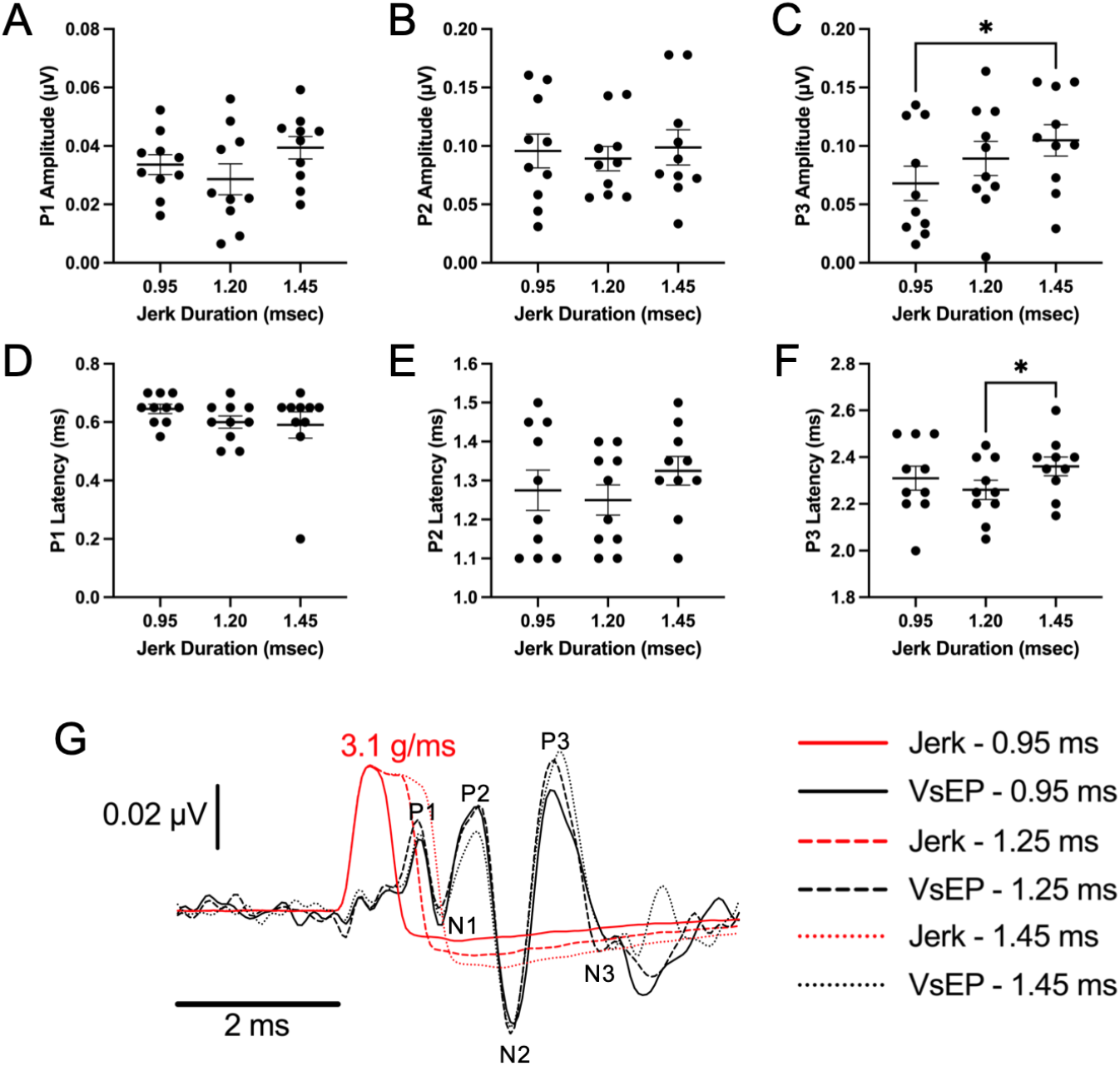
VsEP amplitudes and latencies were compared for P1, P2, and P3 across different jerk durations. Peak amplitudes for P1 **(A)**, P2 **(B)**, and P3 **(C)** are shown for jerk durations of 0.95, 1.20, and 1.45 ms delivered at a constant intensity of 3.1 g/ms. Corresponding peak latencies for P1 **(D)**, P2 **(E)**, and P3 **(F)** are shown across the same jerk durations. Each data point represents an individual animal. Horizontal bars indicate the mean. Error bars represent SEM. Averaged VsEP waveforms recorded at 3.1 g/ms for each jerk duration are shown (G). The jerk stimulus are shown in red (solid = 0.95 ms, dashed = 1.20 ms, dotted = 1.45 ms), and the corresponding VsEP traces are shown in black. Vertical scale bar in G represents VsEP voltage; horizontal scale bar in G represents time. *p < 0.05

In contrast, the P3 component, representing higher-order central vestibular processing, exhibited significant duration-dependent plasticity. The amplitude of the P3 peak was significantly increased in response to the longest tested duration (1.45 ms) compared to the shortest duration (0.95 ms) (Fig. 2C, p < 0.0486). Furthermore, this increase in response magnitude was accompanied by a temporal shift; the P3 latency was significantly delayed at 1.45 ms compared to the 1.20 ms duration (Fig. 2F, p = 0.0320). These findings suggest that while the initial afferent volley is reliably triggered by the leading edge of the kinematic stress, later stages of the central vestibular network integrate stimulus energy over a broadened temporal window, effectively recruiting additional neuronal populations to process sustained inertial inputs.

### 3.2. Peripheral saturation and central recruitment define the dynamic response to stimulus intensity

Having established the network’s sensitivity to stimulus duration, we next evaluated the dynamic range of peripheral and central vestibular populations across increasing jerk intensities (0.64, 3.3, 5.2, and 8.8 g/ms). The VsEP waveforms revealed a stark divergence in how distinct anatomical tiers accommodate escalating inertial stress (Fig 3).

**Figure 3.**
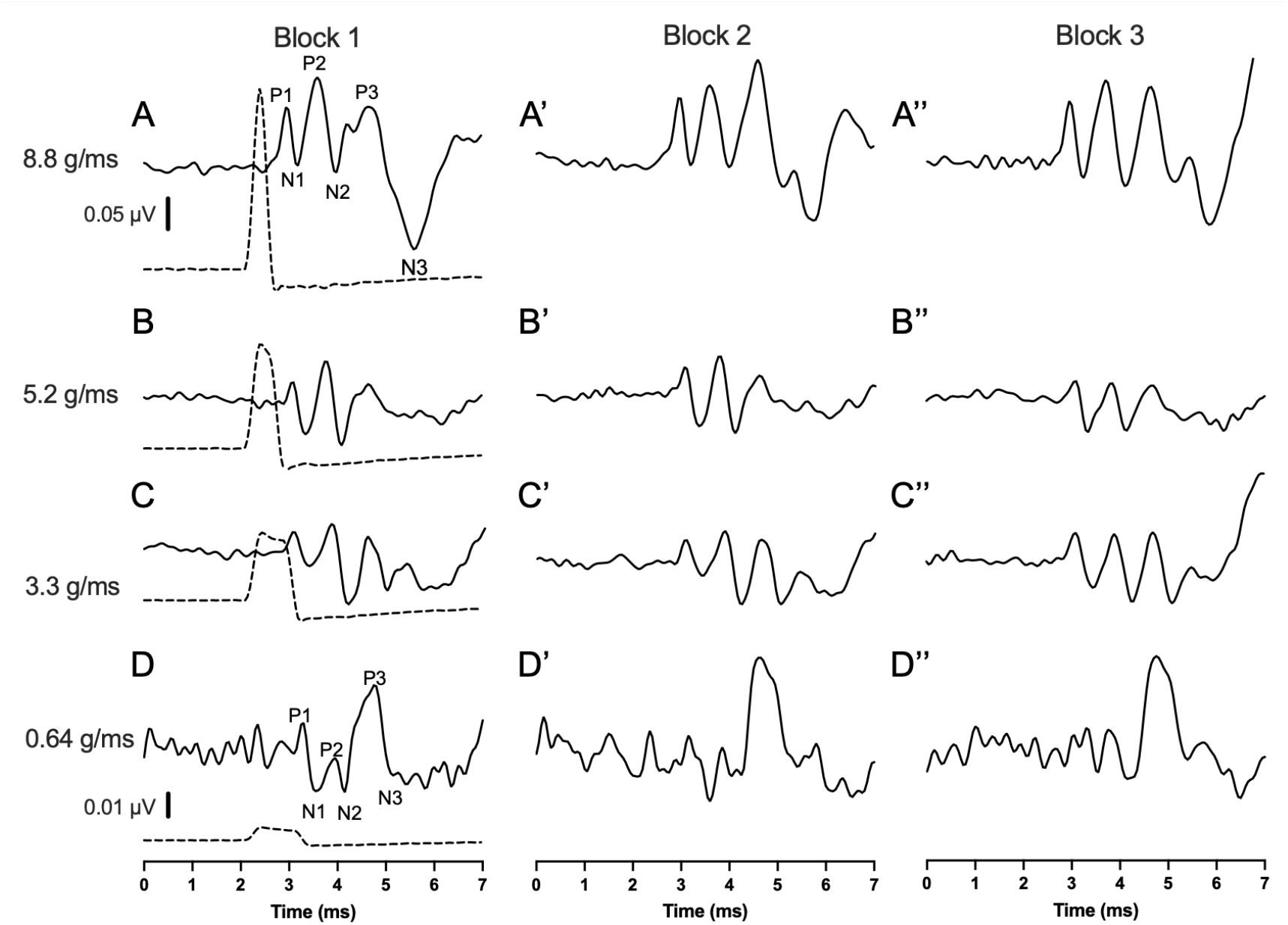
VsEP waveforms represent averaged responses for each jerk intensity within each block of 2,000 jerks. Solid line in all panels represent VsEP trace from a representative animal for each jerk intensity group. Rows depict jerk intensities: 8.8 g/ms **(A– A″)**, 5.2 g/ms **(B–B″)**, 3.3 g/ms **(C–C″)**, and 0.64 g/ms **(D–D″)**. In Block 1 (A–D), the dashed line indicates the jerk stimulus. VsEP peaks (P1, N1, P2, N2, P3, N3) are labeled in A and D. Vertical scale bars represent VsEP voltage: 0.05 µV for A–C and 0.01 µV for D.

Within the peripheral compartment, the P1 amplitude scaled linearly with initial intensity increases; the 0.64 g/ms condition produced significantly smaller amplitudes than the 8.8 g/ms condition (Fig. 4A, p = 0.0197). However, the peripheral response rapidly reached a physiological ceiling, saturating as the stimulus approached 8.8 g/ms, indicating a finite limit to the recruitable pool of striolar irregular afferents. On the other hand, the peak latency of P1 wave showed a larger dynamic range of intensity than P1 amplitude (Fig. 4D). This indicates that the temporal information by P1 nerve activity carries more information about jerk intensity than their aggregated activity level.

**Figure 4.**
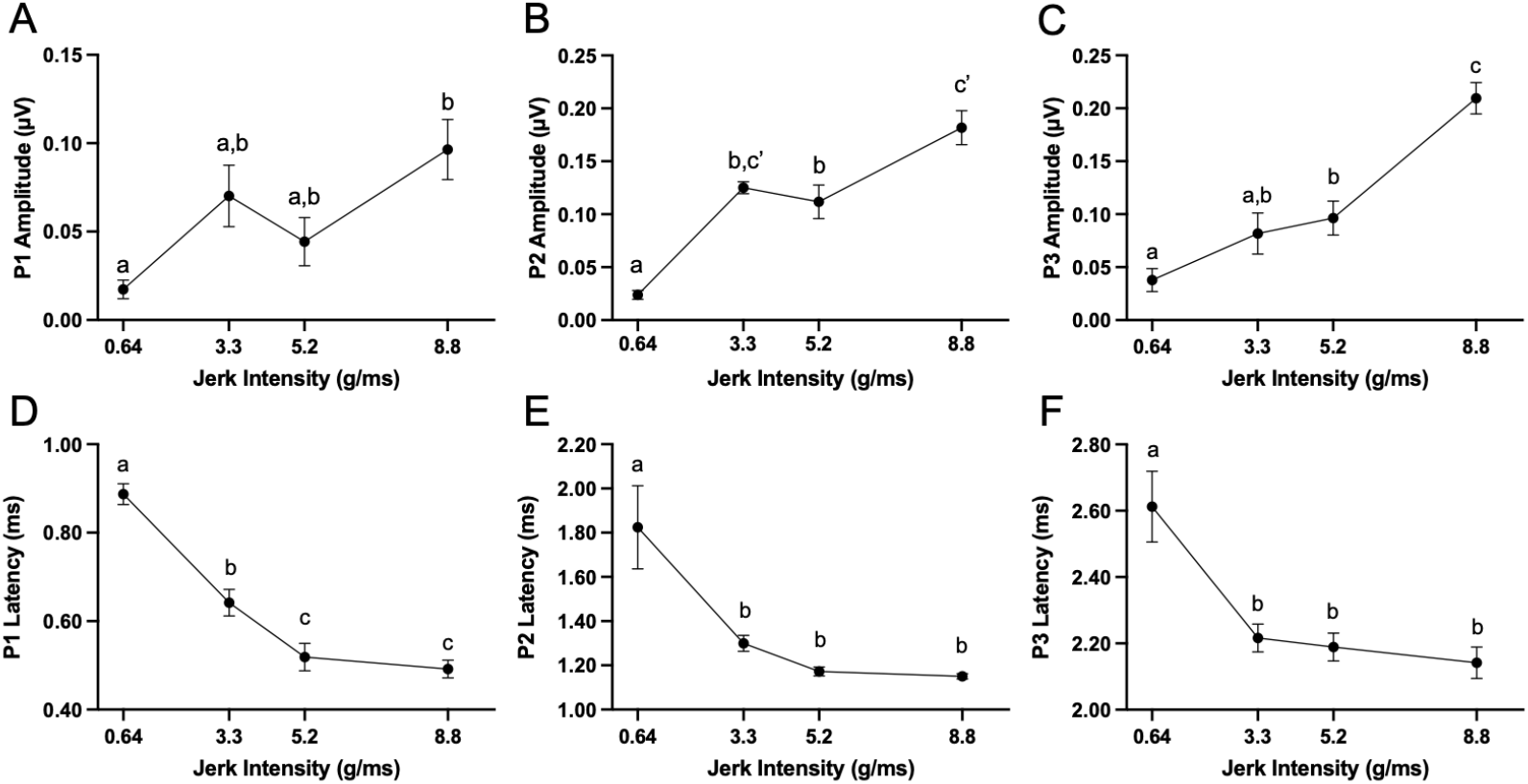
VsEP amplitudes and latencies were compared across different jerk intensities within block 1. Mean peak amplitudes for P1N1 (**A**), P2N2 (**B**), and P3AN3B (**C**) response components are plotted as a function of jerk intensity (0.64, 3.3, 5.2, and 8.8 g/ms). Corresponding response latencies for P1 (**D**), P2 (**E**), and P3 (**F**) are shown across the same jerk intensities. Data points represent group means; error bars represent standard error. Unique letters denote significant difference among intensities (p < 0.05); shared letters denote no significant difference. ‘: denotes p=0.0564 comparing 3.3 and 8.8 g/ms P2 amplitude.

Conversely, the amplitude of the central components (P2 and P3) demonstrated a remarkably broader dynamic range than P1 amplitude, while the latency encoding by P2 and P3 waves deteriorated compared to that by P1 (Fig. 4E-F). Unconstrained by the saturation limits of the aggregated activity level in the periphery, P2 and P3 amplitudes continued to scale with intensity, resulting in significantly larger response magnitudes at 8.8 g/ms compared to all lower intensities (Fig. 4B-C, p < 0.0001 for both P2 and P3). This divergence highlights a central network capable of continuous recruitment. It also points to the shift from the time-based encoding scheme to the amplitude-based one as ascending the vestibular processing pathways. The temporal summation of the peripheral activity by the central regions allows to process extreme inertial forces that overwhelm peripheral capacity in amplitude-encoding of jerk intensity.

### 3.3. Repetitive kinematic stress induces progressive central latency delays

To evaluate how prolonged inertial loading impacts the temporal processing velocity of the vestibular hierarchy, we tracked absolute signal integration times across 6,000 continuous jerk presentations. The application of repetitive stimulation resulted in a systematic, progressive delay in the absolute latencies of the central components of the VsEP waveform. This temporal migration was heavily dependent on the driving force of the stimulus, becoming most pronounced at the highest tested intensity of 8.8 g/ms.

Specifically, significant latency delays accumulated across successive stimulus blocks for the N1, P2, N2, P3A, and N3B peaks (Fig. 5B-I). While the initial peripheral activation timeline remained tightly tethered to its baseline velocity (Fig. 5A), subsequent central processing nodes demonstrated an expanding temporal footprint. This progressive deceleration across the central relays suggests that high-repetition kinematic stress imposes an acute metabolic or vesicle-depleting burden on the quantal synapses of the vestibular nuclei, systematically lengthening the time required for post-synaptic populations to reach depolarization thresholds.

**Figure 5.**
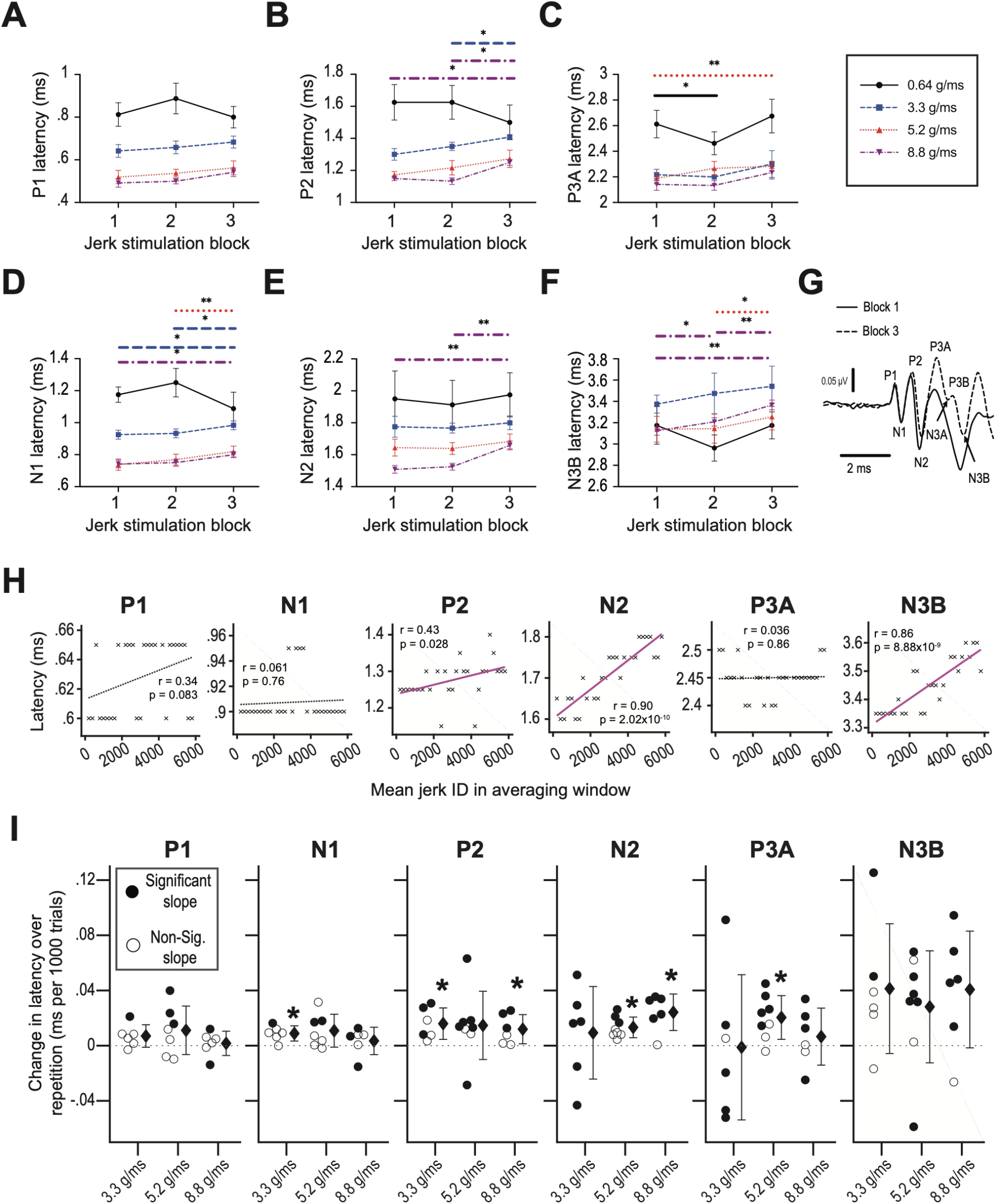
VsEP latencies were compared across repeated blocks and trials at different jerk intensities. Peak latencies for P1 (**A**), P2 (**B**), P3A (**C**), N1 (**D**), N2 (**E**), N3B (**F**) were measured from responses averaged over 2,000 jerks per block (blocks 1–3) in animals stimulated at 0.64, 3.3, 5.2, or 8.8 g/ms. Points denote group mean; error bars represent standard error. Significance bar patterns and colors correspond to the intensity groups indicated in the legend (p < 0.05; Two-way ANOVA; Tukey’s multiple comparisons post-hoc test). Average waveform from the 8.8 g/ms group illustrates peak nomenclature (**G**). Trial-by-trial latency changes were assessed in a representative animal by averaging responses over 400 jerks (positive and negative), with the averaging window advanced in 200-jerk increments within each block, resulting in total 9 x 3 = 27 windows. Latencies were plotted against mean jerk ID (**H**), with significant (magenta) and non-significant (dotted) linear fits (Pearson’s correlation test, p < 0.05). Slopes from these fits are summarized across animals for each peak and intensity (**I**). Filled circles indicate animals with significant correlations; open circles indicate non-significant correlations. Diamonds denote group means; error bars represent standard deviation. Group mean slopes were tested against zero (t-test) and black asterisks indicate marginal significance (p < 0.05, uncorrected for multiple comparisons).

### 3.4. Population amplitudes and initial slope trajectories remain stable across high-repetition blocks

Given the systematic migration of central response latencies, we next examined whether this temporal delay was accompanied by a corresponding decay in overall signal magnitude, which would indicate traditional synaptic fatigue or failure. Surprisingly, analysis of the population data revealed that prolonged repetition did not significantly suppress the overall mean amplitudes of the primary VsEP waves (Fig. 6A-C). The peak-to-peak amplitudes of the P1-N1, P2-N2, and P3A-N3B components remained statistically stable across the entire 6,000-jerk paradigm, with group mean slopes showing no significant deviation from zero (Fig. 6D).

**Figure 6.**
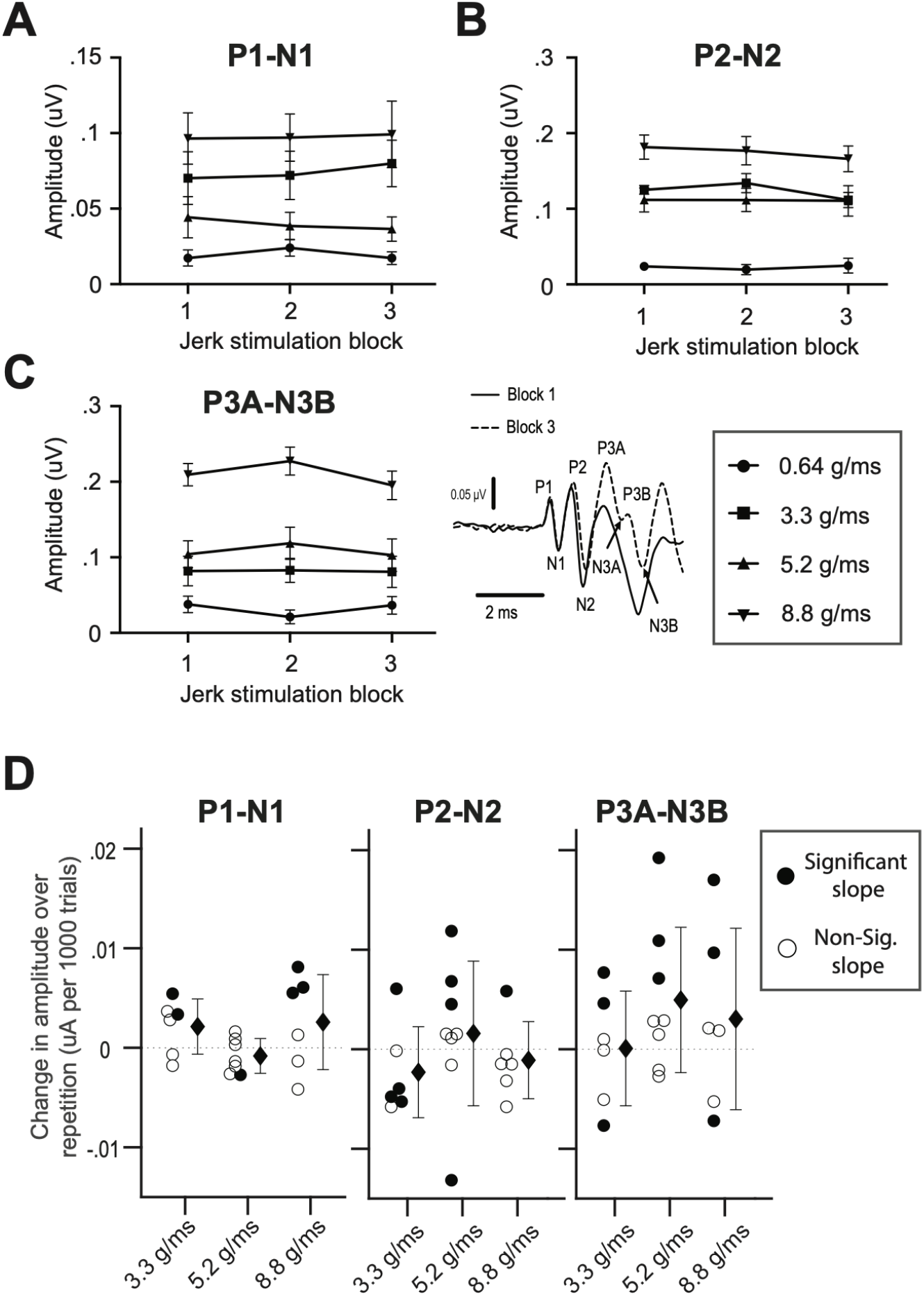
VsEP amplitudes were compared across repeated blocks and trials at different jerk intensities. Peak amplitudes for P1-N1 (**A**), P2-N2 (**B**), and P3A-N3B (**C**) were measured from responses averaged over 2,000 jerks per block (blocks 1–3) in animals stimulated at 0.64, 3.3, 5.2, or 8.8 g/ms. Points denote group mean; error bars represent standard error. No pair of blocks at any jerk intensity showed significant difference in the mean (p > 0.05; Two-way ANOVA; Tukey’s multiple comparisons post-hoc test). Trial-by-trial changes in P1-N1 amplitude (left panel), P2-N2 amplitude (middle panel), and P3A-N3B amplitude (right panel) were assessed across animals by averaging responses over 400 jerks (positive and negative), with the averaging window advanced in 200-jerk increments within each block, resulting in total 9 x 3 = 27 windows (**D**, similar to Fig 5I). Filled circles indicate animals with significant correlations; open circles indicate non-significant correlations. Diamonds denote group means; error bars represent standard deviation. Group mean slopes were tested against zero (t-test).

While the broader population metrics demonstrated macroscopic resilience, fine-grained examination of individual animal trajectories uncovered localized variations in early central processing. At the maximum intensity threshold of 8.8 g/ms, a subset of animals exhibited a progressive flattening of the early central upstroke, characterized by a less steep P2-N2 trajectory. In these specific cases, both the minimum and average voltage slopes between the P2 and N2 peaks decreased linearly as a function of repetition (Fig. 7A-D). However, because this flattening was restricted to a subpopulation, the overarching group means did not achieve statistical significance (Fig. 7E-F). This preservation of global population amplitudes alongside shifting latency profiles suggests that the central network does not simply experience broad failure; rather, it reorganizes its response architecture under sustained loads.

**Figure 7.**
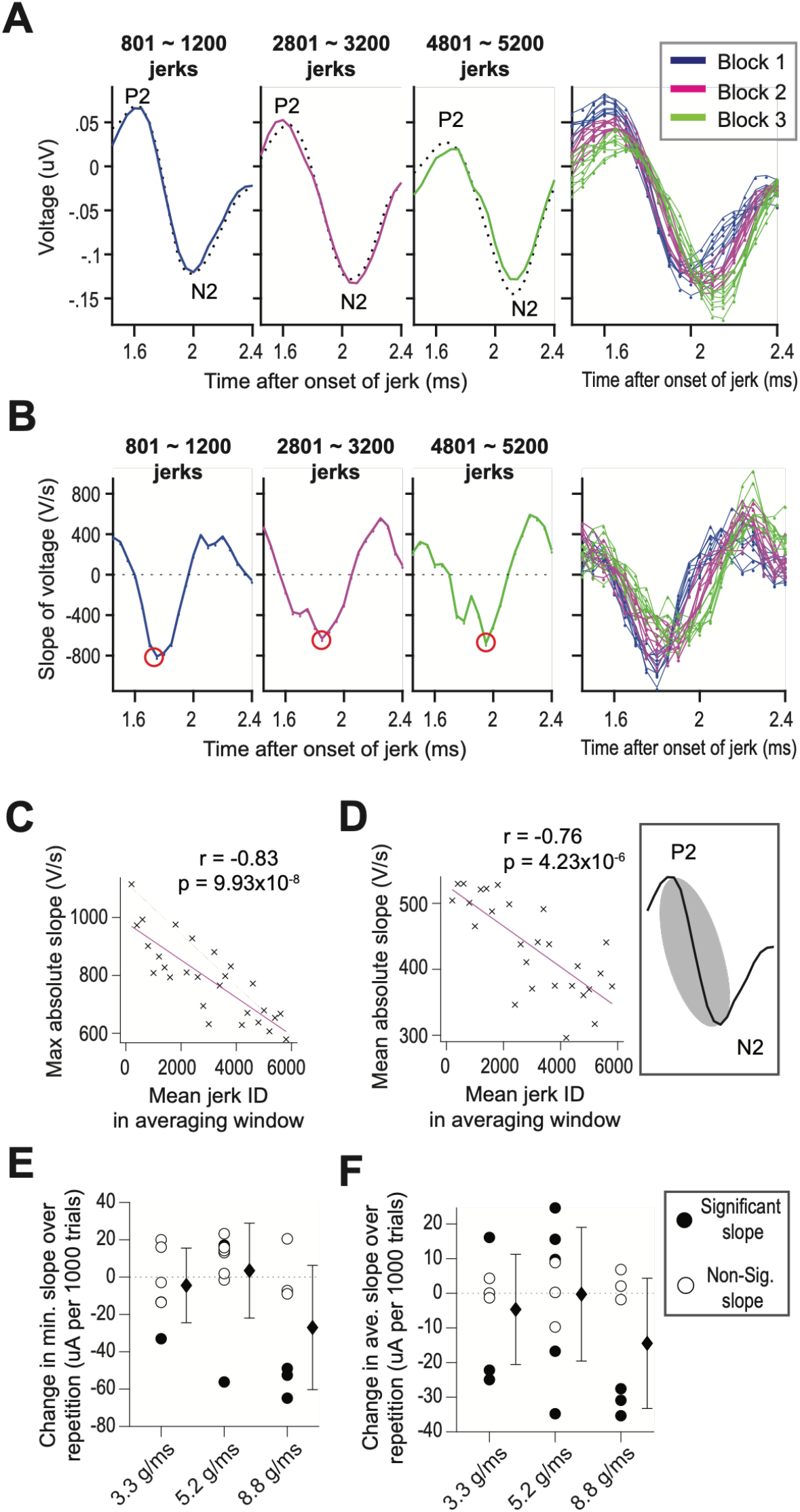
P2N2 slope was compared across repeated jerk trials at different intensities. Example VsEP traces obtained by averaging responses to jerks 801–1,200, 2,801–3,200, and 4,801–5,200 are shown focusing over the P2–N2 downstroke (**A**). Dotted lines show VsEP traces obtained by averaging 2,000 jerk responses in blocks 1, 2, and 3. Traces from all averaging windows containing 400 jerks were superimposed (right-most panel of **A**; blue = block 1, magenta = block 2, green = block 3). The slope (first derivative) of the averaged VsEP traces is shown below each trace, with red circles indicating the most negative slope (steepest downstroke). Slopes of VsEP traces from all averaging windows are also superimposed (right-most panel; **B**). Maximum of the absolute value of slope on the P2–N2 downstroke was plotted against mean jerk ID of averaging windows for a representative animal (Pearson correlation test; **C**), with the significant linear fit shown in magenta. The mean of the absolute value of P2N2 slope (averaged across shaded region in the inset) was similarly plotted against mean jerk ID for the same animal (Pearson correlation; **D**). The estimated slope parameter from the linear fit on max-absolute-slope vs. jerk repetition plots are shown for all animals tested at 3.3, 5.2, and 8.8 g/ms (**E**). Filled circles indicate animals with significant correlations; open circles indicate non-significant correlations. Diamonds denote group means; error bars represent standard deviation. Group mean estimated slopes were tested against zero. The same format is used for the estimated slope of linear fit on average-P2N2-slope vs. jerk repetition plots across the same stimulus intensities (**F**), with group mean slopes tested against zero.

### 3.5. Structural remodeling and steepening of the late-stage central upstroke

As the stimulation blocks progressed, the morphology of the late-stage central VsEP waveform underwent a distinct structural reorganization. Rather than displaying the erratic or degraded profiles typical of a failing sensory system, the late-stage central trace exhibited a progressive smoothing effect. Pre-existing micro-fluctuations, such as early “dips” or “shoulders” embedded within the ascending limbs of the late waves, became less prominent and were systematically absorbed into the broader macro-waveform with continued repetitions (Fig. 8A-F).

**Figure 8.**
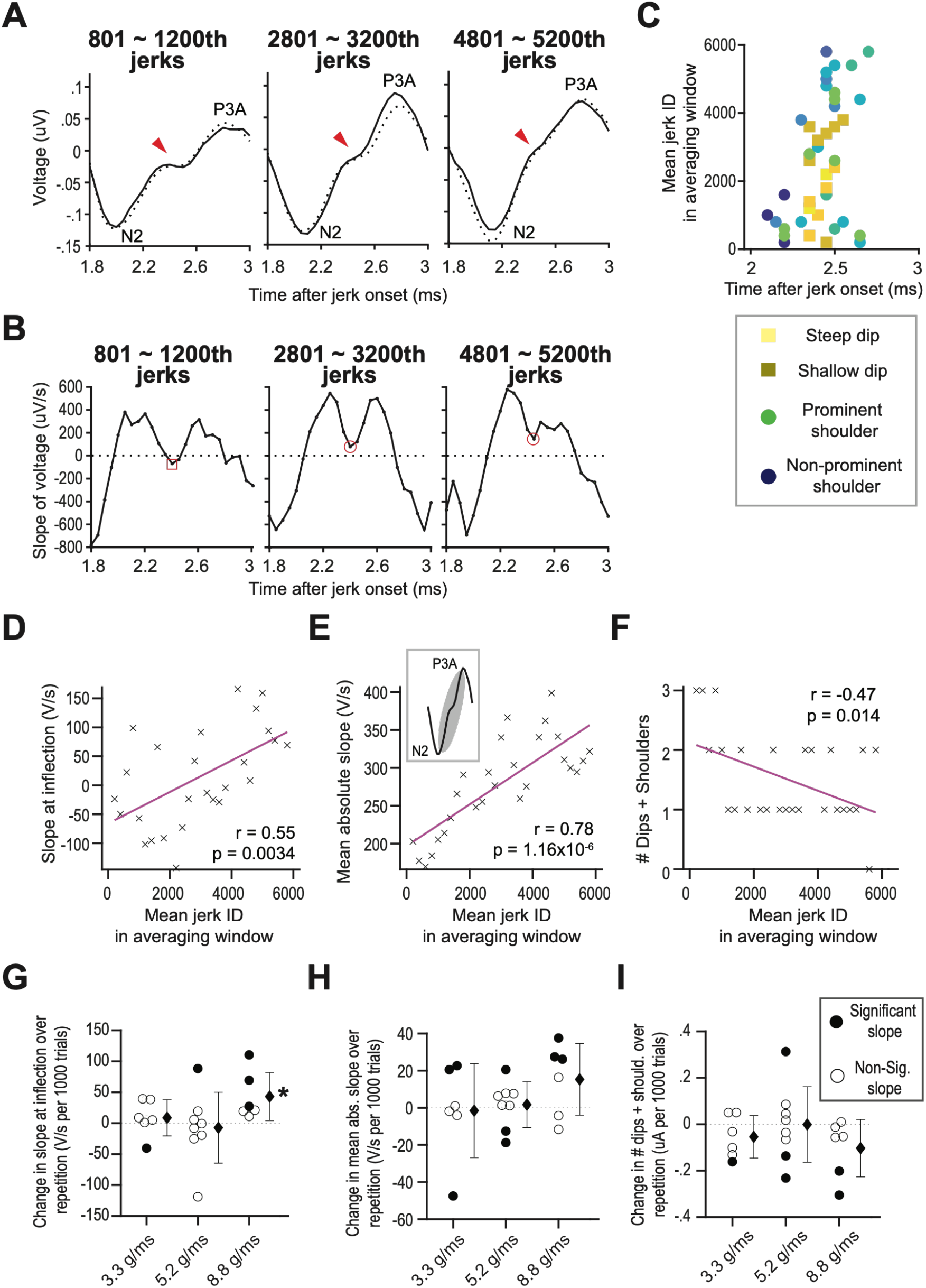
Inflection point and N2–P3A slope were compared across repeated jerk trials at different intensities. VsEP traces were obtained by averaging responses to jerks 801–1,200, 2,801–3,200, and 4,801–5,200, and were analyzed over the N2–P3A upstroke (**A**). Dotted lines show block-averaged VsEP traces for blocks 1, 2, and 3. Traces from all averaging windows were superimposed (left to right panels). Red arrows indicate inflection points corresponding to “dip” and “shoulder” features. The first derivative of the corresponding VsEP traces is shown below each trace (**B**). Red squares indicate local minima crossing below zero, corresponding to a “dip,” where the slope briefly becomes negative. Red circles indicate local minima that do not cross below zero, corresponding to a “shoulder,” where the slope becomes shallower but remains positive. Slope at the inflection point (local minima of the first derivative) was plotted as a function of time of occurrence and mean jerk ID of the averaging windows (**C**). Slope values are color-coded (yellow = more negative, blue = more positive) and symbol-coded (squares = slope < 0; circles = slope > 0). Correlations between mean jerk input and waveform measures are shown for a representative animal, including slope at the inflection point (**D**), mean absolute N2–P3A slope (**E**), and number of dips-plus-shoulders (**F**). When multiple inflection points were present within a single N2–P3A upstroke, the most negative slope was used for that averaging window. Significant linear fits are shown in magenta (Pearson correlation, p < 0.05). Linear fit slopes for all animals tested at 3.3, 5.2, and 8.8 g/ms are shown versus jerk repetition for slope at inflection point (**G**), mean absolute N2–P3A slope (**H**), and number of dips-plus-shoulders (**I**). Filled circles indicate animals with significant correlations; open circles indicate non-significant correlations. Diamonds denote group means; error bars represent standard deviation. Group mean slopes were tested against zero (t-test), with black asterisks indicating marginal significance (p < 0.05, uncorrected for multiple comparisons).

This morphological remodeling was driven by a significant alteration in voltage kinetics between the primary central peaks. The slope at inflection of the voltage trace spanning the interval from the N2 valley to the P3A peak increased significantly at 8.8 g/ms (Fig. 8G). This progressive steepening of the N2-P3A upstroke demonstrates that despite the absolute latency delays introduced by repetitive stress, the rate of late-stage central voltage transformation was actively accelerated. This acceleration provides direct evidence of enhanced postsynaptic synchronization among higher-order central vectors.

### 3.6. Experience-dependent emergence of the synchronized N3A-P3B oscillatory dip

The most striking evidence of central structural plasticity was the robust, experience-dependent emergence of a novel late-stage waveform component. With continued repetition of high-intensity jerk parameters, a secondary, highly synchronized oscillation developed immediately following the P3A peak, designated as the N3A-P3B dip (Fig. 9B). This morphological feature was virtually absent or poorly defined during the baseline and early blocks of stimulation.

**Figure 9.**
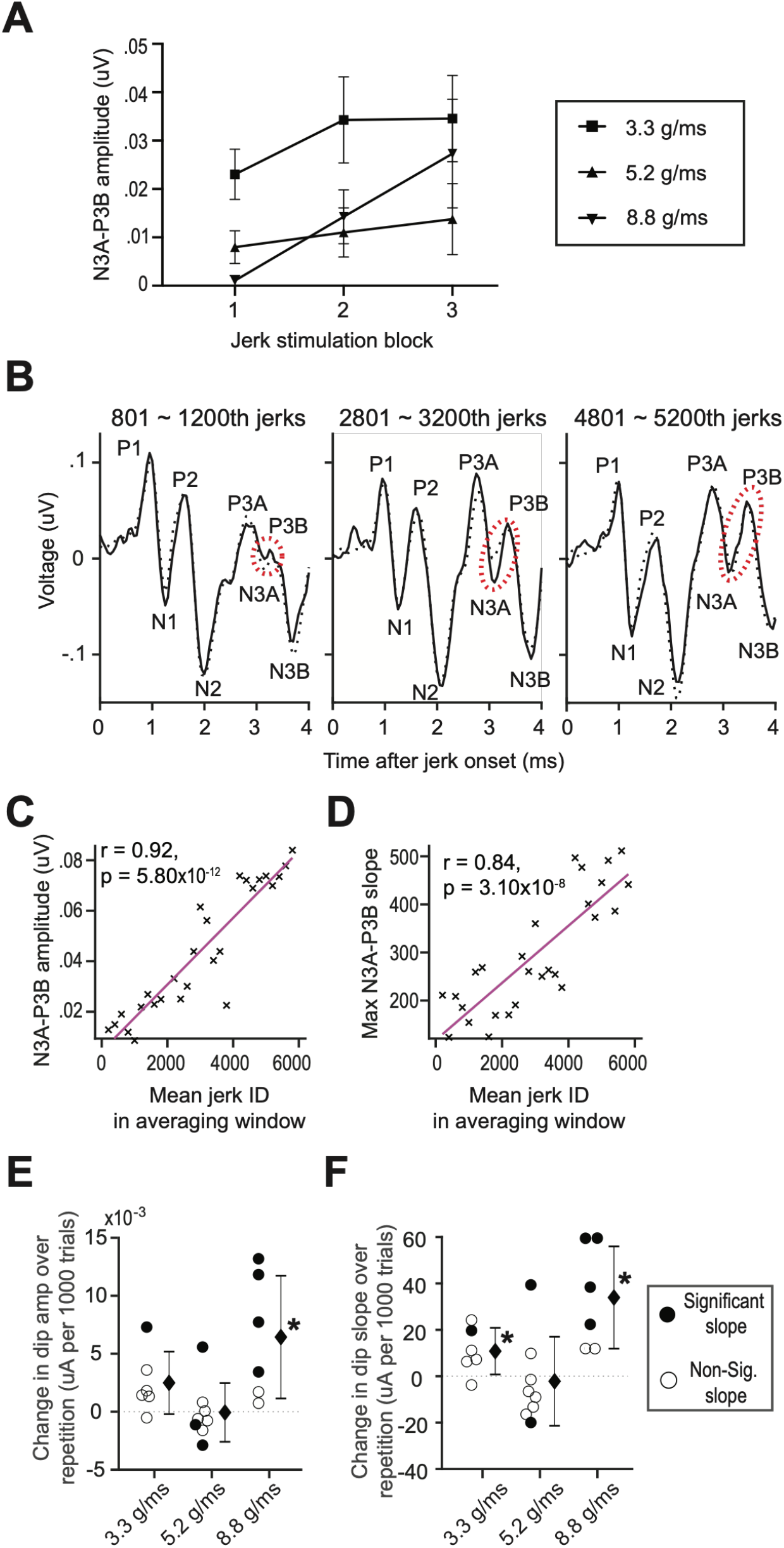
N3A–P3B amplitude and slope were compared across repeated blocks and trials at different intensities. N3A–P3B amplitude was measured on VsEP traces obtained by averaging responses to jerks 1–2,000 in blocks 1, 2, and 3. Three groups of animals were tested at 3.3, 5.2, and 8.8 g/ms. Group mean N3A–P3B amplitudes for blocks 1, 2, and 3 are shown with error bars representing standard error (**A**). VsEP traces were obtained by averaging responses to jerks 801–1,200, 2,801–3,200, and 4,801–5,200 for a representative animal (left to right panels, **B**). Dotted ellipses indicate the increasing N3A–P3B “dip” across repeated jerk presentations. N3A–P3B amplitude (**C**) and maximum slope (**D**) were measured in 400-jerk windows and plotted against mean jerk ID of the averaging windows. Significant linear fits are shown in magenta (Pearson correlation, p < 0.05). Changes in N3A–P3B “dip” amplitude (**E**) and slope (**F**) are shown for all animals tested at 3.3, 5.2, and 8.8 g/ms. Filled circles indicate animals with significant correlations; open circles indicate non-significant correlations. Filled diamonds denote group means, with error bars representing standard deviation. Group mean slopes were tested against zero (t-test). Black asterisks indicate marginal significance (p < 0.05, uncorrected for multiple comparisons).

As the tracking paradigm continued, both the absolute amplitude and the maximum descending slope of this N3A-P3B dip increased significantly across successive stimulation blocks (Fig. 9C-F). This structural phenotype was highly state-dependent, surfacing prominently when driven by the extreme inertial stress of the 8.8 g/ms threshold. The progressive crystallization of this oscillatory dip suggests that the central nervous system, when confronted with an unremitting stream of saturated peripheral inputs, adaptively recruits or unmasks a downstream, resonant neuronal population that fires with increasing tight temporal alignment.

### 3.7. Phase coherence and time-frequency dynamics of central neural honing

To uncover the underlying biophysical mechanism driving the morphological remodeling and the emergence of the N3A-P3B oscillation, the single-trial electrophysiological data was subjected to CWT and ITPC analyses (Fig. 10). This time-frequency approach allowed us to move beyond static, averaged waveforms and evaluate the real-time phase alignment of the underlying sensory networks across all 6,000 trials.

**Figure 10.**
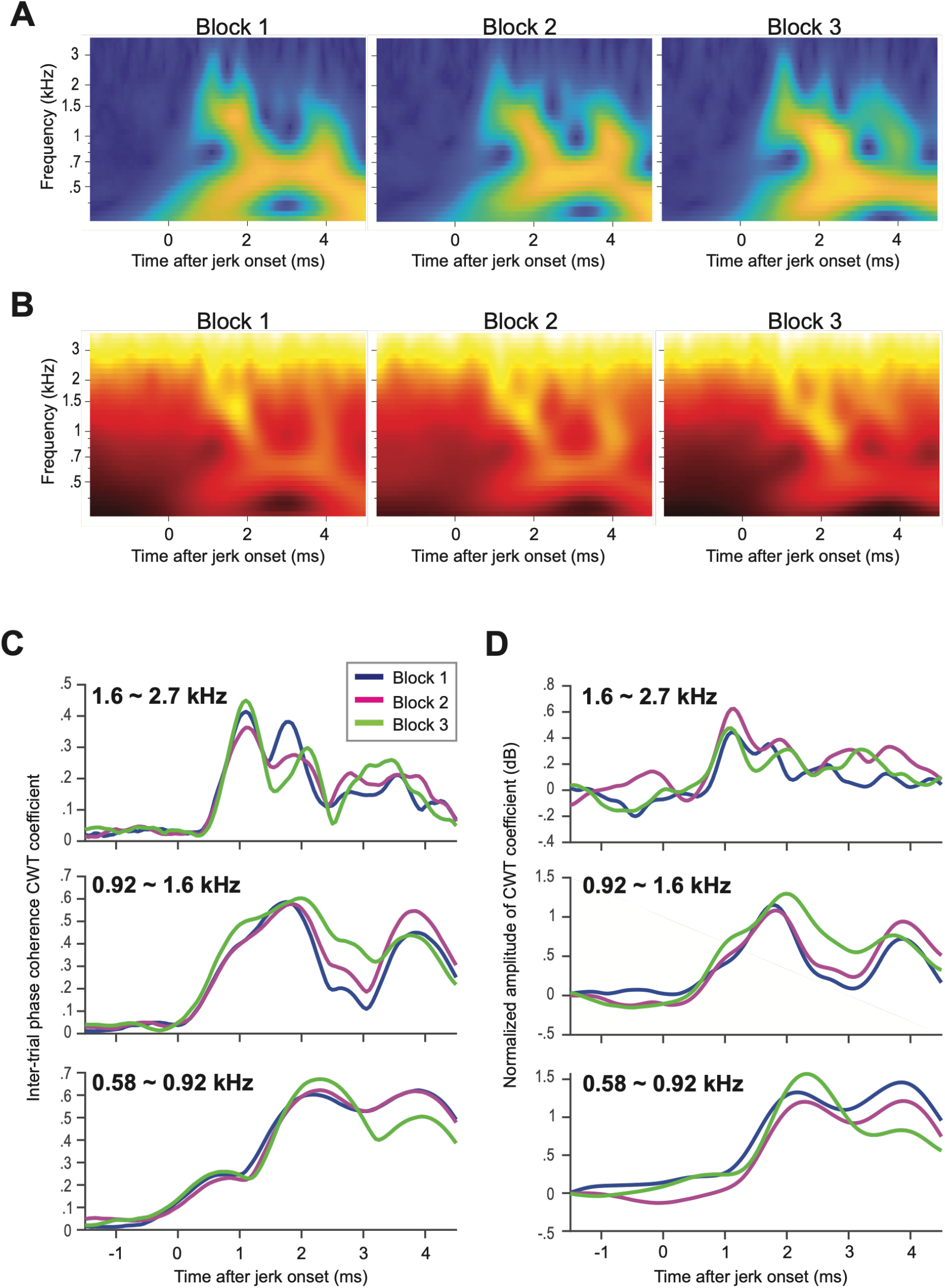
Continuous Wavelet Transform (CWT) was performed on VsEP response to a single pair of positive and negative jerks. Inter-trial-phase-coherence (ITPC) of WT coefficients is shown as a two-dimensional (Time x Frequency) heat map for block 1, 2, and 3 (left to right, **A**). The amplitude of the CWT coefficients is shown as a two-dimensional (Time x Frequency) heat map for block 1, 2, and 3 (left to right, **B**). ITPC of CWT coefficients is averaged across 0.58∼0.92 kHz (bottom panel), 0.92 ∼1.6 kHz (middle panel), and 1.6∼2.7kHz (top panel), and plotted as a function of time to compare between block 1, 2, and 3 (**C**). Normalized amplitude of CWT coefficients is averaged across 0.58∼0.92 kHz (bottom panel), 0.92 ∼1.6 kHz (middle panel), and 1.6∼2.7kHz (top panel), and plotted as a function of time to compare between block 1, 2, and 3 (**D**).

The mathematical decomposition of the single-trial waveforms revealed that repetitive kinematic stimulation selectively refined the trial-to-trial timing of mid-frequency wave oscillations spanning the 0.92 to 1.6 kHz range (and 1.6 to 2.7 kHz range in some degree). This progressive phase-locking did not occur uniformly across the entire response window; instead, it was heavily concentrated within a post-stimulus temporal window of 2.0 to 3.4 ms. This localized surge in phase coherence aligned with the exact temporal coordinate where the N3A-P3B oscillatory dip emerges in the macro-averages.

By demonstrating that the trial-to-trial jitter of these mid-frequency waves is systematically minimized as repetitions increase, the time-frequency data provides definitive proof of a central “neural honing” mechanism. The central network actively compensates for early synaptic delays by tightening the phase-coherence of its downstream targets, ensuring the maintenance of high-fidelity synchronous throughput under continuous mechanical stress.

In contrast, there is a localized depression in phase coherence spanning 1.6 to 2.7 kHz within a post-stimulus temporal window of 1.5 to 2.0 ms. This corresponds to the previously described progressive flattening of the early central upstroke, characterized by a less steep P2-N2 trajectory. This may be reflecting the metabolic and vesicular depletion inherent to highly active quantal synapses in the early VNC processes.

## 4. DISCUSSION

The hierarchical response of the mammalian vestibular system to linear jerk stimulation represents a highly adaptive, multi-tiered biological signal processing network. Our findings establish a fundamental dichotomy between the stable, fatigue-resistant sensory periphery and the highly plastic, state-dependent central vestibular pathways. By mapping the physiological boundaries of this system up to extreme intensities (8.8 g/ms) and under prolonged repetitive stress, the present work demonstrates that while the amplitude-encoding by the peripheral reaches a hard biological ceiling, central networks engage in a dynamic process of short-term synaptic plasticity and “neural honing” to preserve critical homeostatic throughput.

### 4.1. Peripheral Stability and the Limits of Non-Quantal Transmission

The robust stability of the initial afferent volley, represented by the VsEP P1 component, underscores the specialized architecture of the vestibular periphery as a rapid transient detector. Our results show that P1 amplitude saturates at high intensities and its latency remains remarkably invariant during repetitive stimulation, indicating a fatigue-resistant sensor and therefore, a secure encoding scheme that the higher processing stations can rely on. This physiological capability is best explained by the unique or specialized synaptic mechanics at the striolar neuroepithelium. Irregular afferent action potentials are heavily dependent on non-quantal synaptic transmission between type I hair cells and calyceal afferent terminals [13-17]. This specialized transmission mechanism bypasses the traditional vesicular release cycle, eliminating synaptic delay and allowing for sub-millisecond temporal fidelity and precise latency encoding. However, this speed comes at the cost of dynamic range in the amplitude-based encoding; the system reaches its maximum recruitment limit relatively early, completely synchronizing the available pool of striolar afferents and resulting in the observed saturation of the P1 amplitude.

### 4.2. Central Recruitment and State-Dependent Quantal Summation

In stark contrast to the periphery, the central vestibular components (P2 and P3) demonstrate a vastly superior dynamic range, continuing to scale in amplitude even after the peripheral signal has saturated. However, in parallel, the latency encoding by P2 and P3 waves deteriorated compared to that by P1. This divergence highlights a fundamental shift from the temporal-to rate- (amplitude-) coding which is likely the manifestation of the shift from non-quantal signal propagation to state-dependent quantal summation. Because the P2 and P3 waves are generated through classical quantal transmission from irregular afferents to second-order neurons within the vestibular nuclear complex (VNC) and subsequent central vestibular nuclei, they are subject to temporal and spatial summation [18]. This classical synaptic architecture allows the central pathways to pool multiple inputs over slightly broadened integration windows, effectively recruiting additional neuronal populations that possess higher activation thresholds [19]. This mechanism of central recruitment explains how the vestibular network successfully processes and scales its responses to intense inertial forces that overwhelm the overall activity level of the peripheral sensory apparatus.

### 4.3. Adaptive Plasticity and Central “Neural Honing”

The application of intense, repetitive linear jerk (up to 6,000 continuous stimuli) revealed a profound adaptive capability within these central, quantal networks. While repetitive stimulation typically induces synaptic fatigue and uniformly degrades sensory signaling, our trial-by-trial CWT analysis revealed a more sophisticated compensatory mechanism. It is true that repetitive stress systematically delayed absolute signal integration times for central peaks (e.g., N1, P2, N2, P3A), likely reflecting the metabolic and vesicular depletion inherent to highly active quantal synapses. This degradation was evident in some animals who showed the P2 slope shallowing with jerk repetition. According to the wavelet analysis, this is likely due to degradation of Inter-trial phase coherence in the high-frequency VsEP components. However, rather than simply degrading, the central network actively shifted its functional focus.

This shift is characterized by the robust emergence of a late-stage oscillation, the N3A-P3B component, whose amplitude and steepness significantly increased with continued stimulation. ITPC analysis confirms that this emergence is driven by a significant improvement in the trial-to-trial timing and absolute strength of mid-frequency wave oscillations (0.92–1.6 kHz) underlying the late VsEP response. We term this phenomenon “neural honing.” As the earlier central synapses (P2) exhibit signs of delayed propagation, the system undergoes short-term synaptic plasticity, adaptively tightening the synchrony of downstream vestibulocerebellar or higher-order vestibular integrations.

### 4.4. Study Limitations and Future Applications

The present study was not designed to examine sex as a biological variable, providing opportunities for future studies to consider sex differences. Additionally, our examination of varied jerk intensity (Fig. 4) used between-animal comparisons, as VsEP responses exhibit greater variability across animals than within the same animal [20]. Paired intensity comparisons in future studies may further resolve within-animal jerk intensity response dynamics. Finally, the current study was focused on physiological analyses and did not include histological analyses to directly assess the synaptic and activity-related mechanisms associated with the observed VsEP changes. Future histological studies will provide additional insight into these neurological mechanisms.

The findings of our study are directly applicable in the design of future experiments using VsEP to investigate vestibular hypofunction following vestibular trauma, including ototoxicity, aging, noise exposure, and blast overpressure. Since our results indicate that repetition and large intensities confound with VsEP responses, future studies should carefully consider these parameters when developing acquisition protocols and avoid excessive repetition. Moreover, studies should assess functional changes in vestibular-related motor performance following kinematic stress and whether temporal dynamic VsEP changes may predict vestibular symptoms including fall risk.

By synthesizing these temporal findings, this study suggests that the central vestibular system is not a passive relay, but an actively gating network that redistributes synchronous firing to maintain sensory fidelity during continuous kinematic stress. This framework provides a critical foundation for diagnosing acceleration-induced vestibular hypofunction and understanding how state-dependent plasticity protects against inertial trauma.

## Acknowledgements

We thank Maryam Abbawi and Chloe Daniels for their contribution to data processing and analysis.

## Funding

US Department of Veterans Affairs grant 1I01RX001986 (A.G.H. and F. Akin)

US Department of Veteran Affairs 1I21RX004111 (A.G.H.)

National Institutes of Health grant R56DC021073 (A.G.H.)

The views expressed do not necessarily reflect the official policies of the Department of Health and Human Services, nor does mention of trade names, commercial practices, or organizations imply endorsement by the US Government.

## Author Contributions

Syed Danial Naqvi (SDN), Mamiko Niwa (MN), Rod D. Braun (RDB), Mirabela Hali (MH), Aaron K. Apawu (AKA), Avril Genene Holt (AGH)

Conceptualization: RDB, AGH

Data curation: SDN, MH, AKA

Formal analysis: SDN, MN, RDB, AGH

Methodology: SDN, RDB, AGH

Investigation: SDN, MH, AKA

Visualization: SDN, MN, RDB, AGH

Resources: SDN, RDB, MH, AGH

Software: SDN, MN, RDB, AGH

Validation: SDN, MN, RDB, AGH

Funding acquisition: AGH

Project administration: RDB, AGH

Supervision: RDB, AGH

Writing—original draft: SDN, MN, RDB, AGH

Writing—review & editing: SDN, MN, RDB, AGH

**All authors declare they have no competing interests**.

## Data availability

The datasets generated during and/or analyzed during the current study are available from the corresponding author on reasonable request.

## Notes

### Competing Interest Statement

The authors have declared no competing interest.

